# Distinct mechanical signatures of dairy and plant-based cheeses

**DOI:** 10.64898/2026.11.05.764742

**Authors:** Aeneas O. Koosis, Ethan C. Darwin, Vilde Ellingsen, Ellen Kuhl

**Affiliations:** Department of Mechanical Engineering, Stanford University, Stanford, CA 94305, USA; Department of Civil and Environmental Engineering, Stanford University, Stanford, CA 94305, USA

## Abstract

Plant-based cheeses account for about 1% of U.S. retail cheese sales and often fail to match the texture and flavor of dairy cheese. Here we compare three commercial dairy and plant-based pairs, cheddar, feta, and mozzarella, with texture profile analysis (*n* = 60), tension, compression, and shear tests (*n* = 219), constitutive artificial neural networks, and a consumer panel (*n* = 306). Plant mozzarella is at least five times stiffer than dairy mozzarella at small strain, yet softer and less cohesive under 50% compression. Plant feta matches the initial stiffness of dairy feta but shows a peak in its shear response. Plant cheddar differs in both stiffness and shear response. Model discovery selects first-invariant models for all six cheeses, but no model term separates the two groups. Consumers rate 68–78% of attributes “just about right” for dairy cheeses and 29–38% for plant-based cheeses. These single-product comparisons define product-specific targets for plant-based cheese design.

## Introduction

Dairy production contributes substantially to the environmental footprint of the food system, and cheese is among the more greenhouse-gas-intensive foods per kilogram ^1–3^. Retailers now routinely stock dairy-free cheese ^4^, and replacing dairy with plant-based products can lower emissions ^5^. The global market for dairy alternatives may reach $66.9 billion by 2030^6^, but plant-based cheese accounts for only about 1% of U.S. retail cheese sales by dollar value^4^. Wider adoption of plant-based cheese depends on consumer acceptance.

In an open-source benchmark adapted from a study of meat^7^, 2,183 consumers evaluated commercial dairy and dairy-free products across ten categories^8^. Dairy-free barista milk, creamer, and fluid milk received sensory ratings comparable to those of their dairy counterparts, but consumers consistently rated dairy-free cheese lower than dairy cheese. Dairy-free mozzarella received liking scores 1.46 points below its dairy benchmark on a nine-point scale, and consumers liked samples with off-flavors, artificial notes, or bitterness less.

Dairy cheese texture arises mainly from the casein network, which retains fat and water and deforms and fractures under load ^9;10^. Many commercial plant-based cheeses instead rely on combinations of starch, plant-derived fats, proteins, and hydrocolloids, and the fat type alone can change their hardness and melt ^11–13^. These products often fail to match the texture and mouthfeel of dairy cheese ^14–17^. Food scientists widely use texture profile analysis (TPA) to characterize cheese texture. TPA measures hardness, cohesiveness, and chewiness in a double compression cycle^18;19^, but it does not independently characterize tensile or shear behavior. Complementary loading modes can reveal differences in stiffness and deformation-dependent response that double compression alone does not resolve.

Constitutive artificial neural networks (CANNs) represent the strain energy as a weighted sum of interpretable functions and select compact mathematical models that best fit the data^20–22^. Applied to tension, compression, and shear tests, this approach has distinguished plant-based from animal meat^23–27^ and has characterized fungal steaks^28;29^ and tofu ^30^. Paired with consumer testing, the same measurements have partly explained how texture shapes consumer acceptance^31;32^. CANNs belong to a wider effort to use artificial intelligence for food design ^33–35^. Together with consumer evaluation, multimodal mechanical testing and interpretable constitutive models can show where individual plant-based cheeses differ from their dairy counterparts.

Here we combine TPA, multimodal mechanical testing, constitutive model discovery, and consumer evaluation to examine how differences between dairy and plant-based cheeses vary across three cheese styles: cheddar, feta, and mozzarella. Through direct comparisons of one commercial dairy and one commercial plant-based product within each style, we assess whether mechanical differences are common to the three pairs or specific to individual products, and how well the discovered models capture those differences. We also compare consumer responses within each pair and explore relationships between TPA parameters and perceived texture across the six products.

## Results

### Study design

We test one commercial dairy and one commercial plant-based product in each of three cheese styles (Table 1). We measure each product with TPA and with tension, compression, and shear tests (Fig. 1), and we fit the tension, compression, and shear data with a constitutive artificial neural network (Fig. 2).

**Figure 1:**
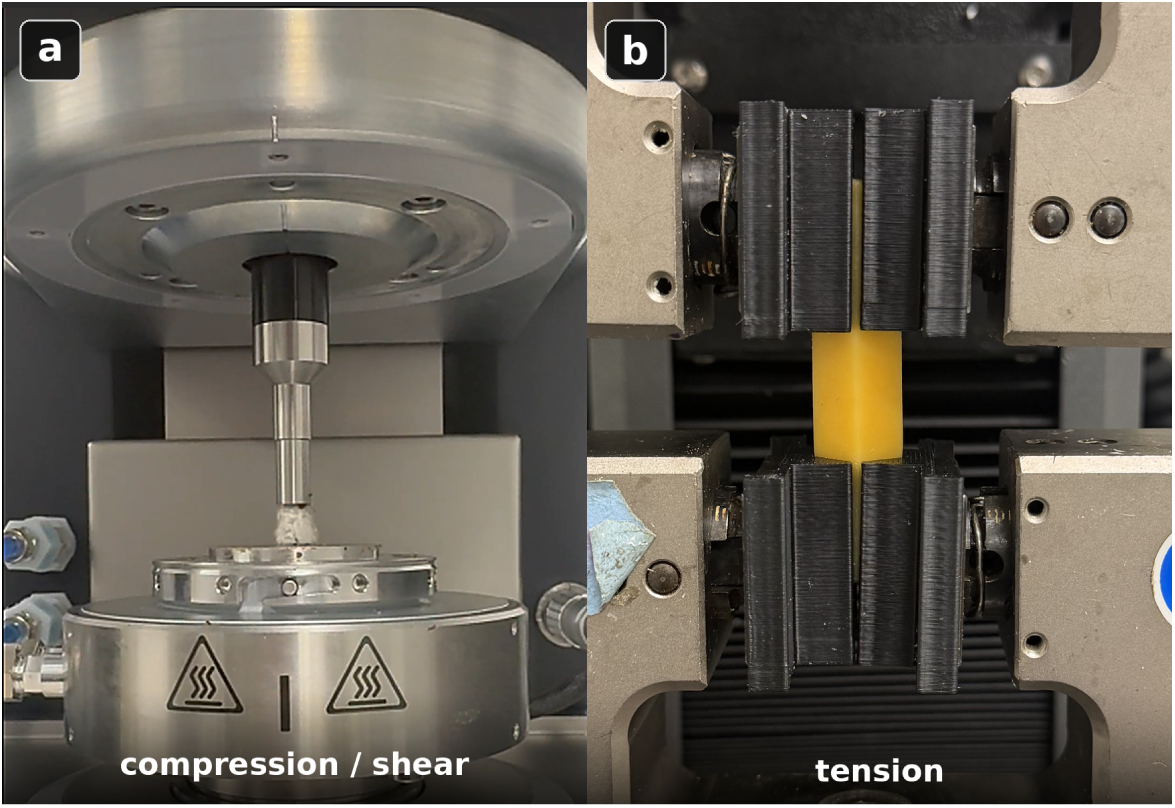
Mechanical test configurations. (a) The rheometer’s parallel plates hold a cheese core for compression and simple shear tests. Sandpaper surfaces prevent slip. (b) The tensile grips hold a cheese specimen between 3D-printed serrated faces.

**Figure 2:**
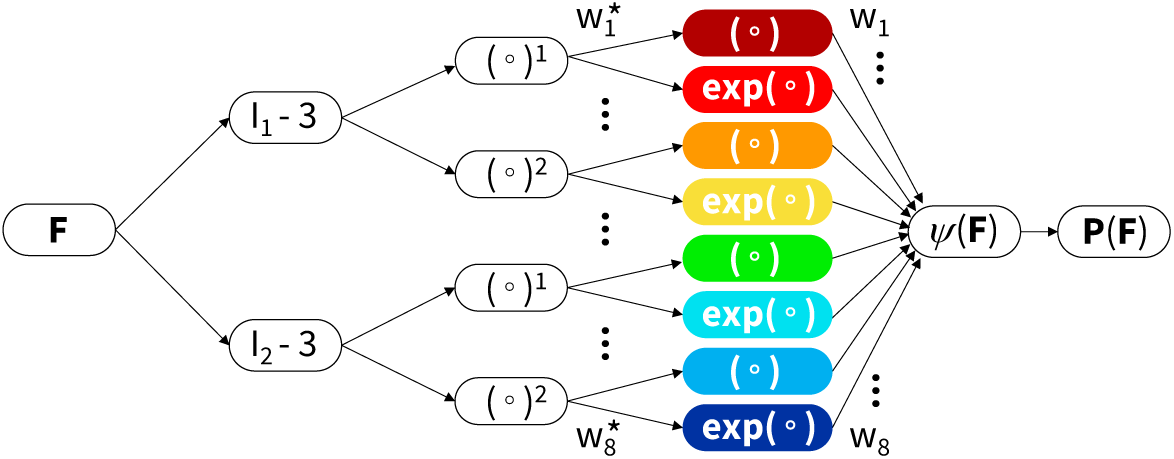
Constitutive artificial neural network. The network takes the deformation gradient ***F*** as input and computes the shifted invariants [*I*_1_ *−* 3] and [*I*_2_ *−* 3]. The first hidden layer raises them to the first and second powers, (*◦*)^1^ and (*◦*)^2^, and the second hidden layer applies the identity (*◦*) and the exponential exp(*◦*) *−* 1. The weighted sum of the eight color-coded terms gives the strain energy function *ψ*, and its derivative, together with the pressure constraint, gives the Piola stress ***P*** .

**Table 1:**
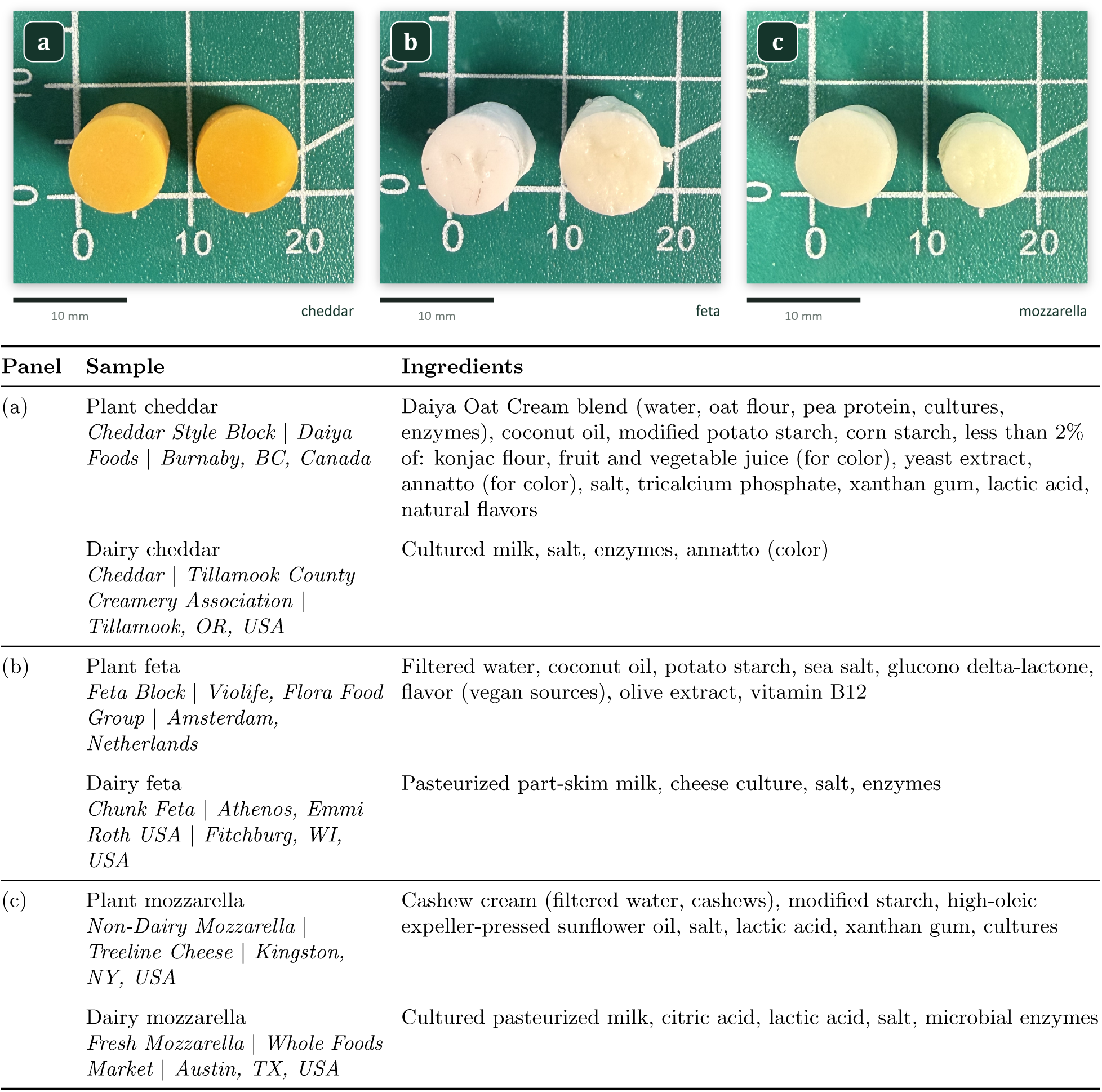
Cheese products and ingredient lists. Photographs show (a) cheddar, (b) feta, and (c) mozzarella cores from above on a 10 mm grid. Each panel shows the plant-based core on the left and the dairy core on the right. Product entries list product name, company, and company location. Ingredient lists follow the product labels.

### Texture profile analysis

Each plant-based cheese differs from its dairy counterpart in a different set of TPA parameters (Table 2; Figs. 3 and 4). Plant cheddar has higher hardness than dairy cheddar (26.2 *±* 1.5 vs. 13.3 *±* 2.5 N, *p <* 0.001), but comparable cohesiveness (*p* = 0.15) and chewiness (*p* = 0.06). Plant feta has lower hardness (5.8 *±* 0.5 vs. 7.6 *±* 1.8 N, *p* = 0.010), cohesiveness (0.08 *±* 0.02 vs. 0.17 *±* 0.03, *p <* 0.001), and chewiness (0.17 *±* 0.14 vs. 0.81 *±* 0.16 N, *p <* 0.001) than dairy feta. Plant mozzarella also has lower hardness (6.3 *±* 0.7 vs. 10.0 *±* 0.9 N), cohesiveness (0.14 *±* 0.06 vs. 0.58 *±* 0.02), and chewiness (0.69 *±* 0.58 vs. 5.13 *±* 0.47 N) than dairy mozzarella (all *p <* 0.001).

**Figure 3:**
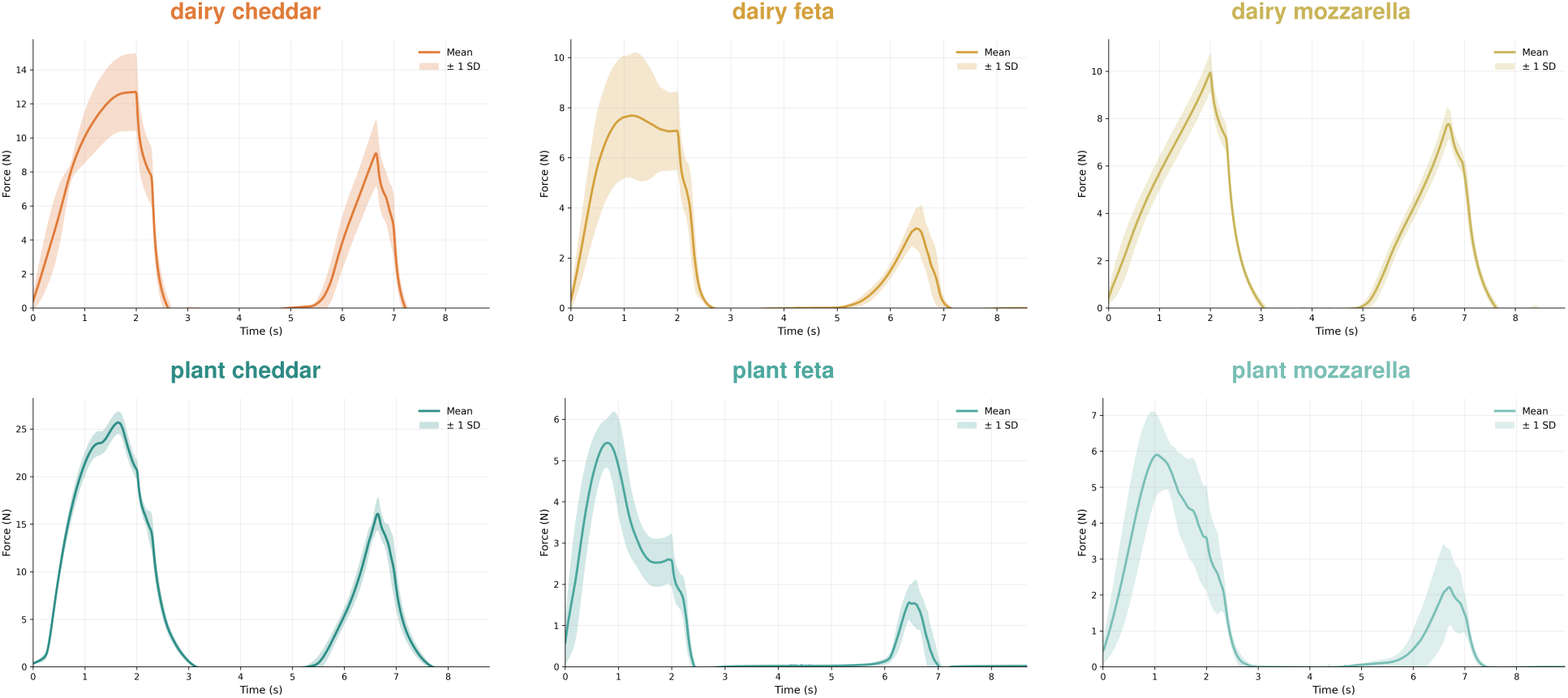
Texture profile analysis: force–time curves. Mean force–time curves during two consecutive compression cycles for all six cheese products, with *n* = 9–11 specimens per product. Shaded bands indicate *±* standard deviation.

**Figure 4:**
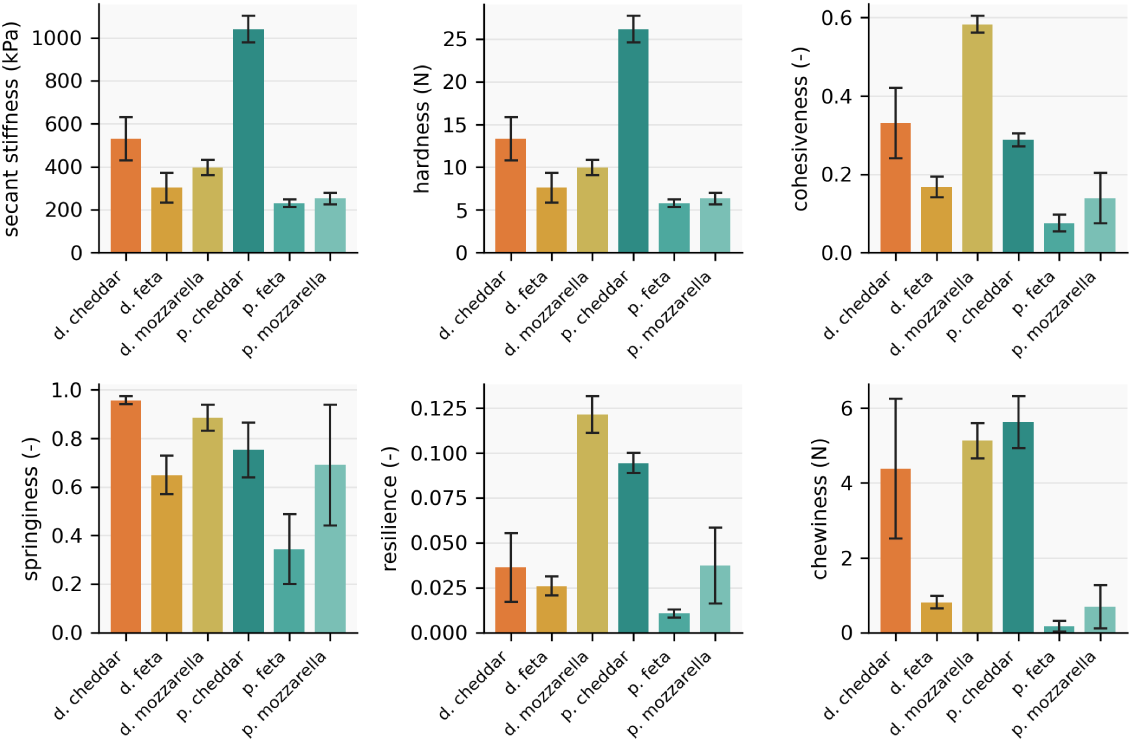
Texture profile analysis: texture parameters. Stiffness index, hardness, cohesiveness, springiness, resilience, and chewiness for all six cheese products, with *n* = 9–11 specimens per product. Error bars indicate *±* standard deviation.

**Table 2:**
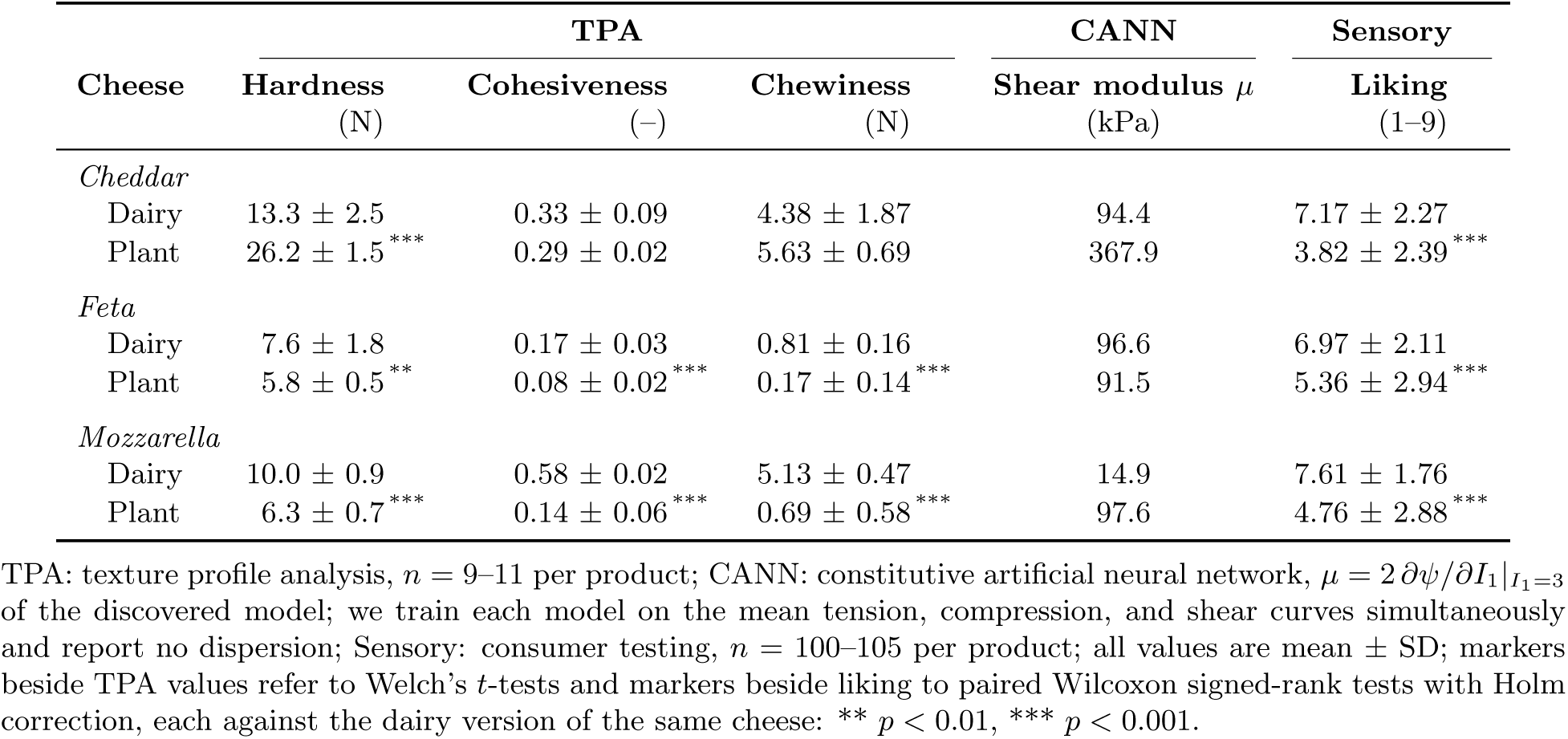
Instrumental texture and consumer liking.

|  | **TPA** |  |  | **CANN** | **Sensory** |
|---|---|---|---|---|---|
| **Cheese** | **Hardness**<br>(N) | **Cohesiveness**<br>(–) | **Chewiness**<br>(N) | **Shear modulus *μ***<br>(kPa) | **Liking**<br>(1–9) |
| *Cheddar* |  |  |  |  |  |
| Dairy | 13.3 ± 2.5 | 0.33 ± 0.09 | 4.38 ± 1.87 | 94.4 | 7.17 ± 2.27 |
| Plant | 26.2 ± 1.5<sup>***</sup> | 0.29 ± 0.02 | 5.63 ± 0.69 | 367.9 | 3.82 ± 2.39<sup>***</sup> |
| *Feta* |  |  |  |  |  |
| Dairy | 7.6 ± 1.8 | 0.17 ± 0.03 | 0.81 ± 0.16 | 96.6 | 6.97 ± 2.11 |
| Plant | 5.8 ± 0.5<sup>**</sup> | 0.08 ± 0.02<sup>***</sup> | 0.17 ± 0.14<sup>***</sup> | 91.5 | 5.36 ± 2.94<sup>***</sup> |
| *Mozzarella* |  |  |  |  |  |
| Dairy | 10.0 ± 0.9 | 0.58 ± 0.02 | 5.13 ± 0.47 | 14.9 | 7.61 ± 1.76 |
| Plant | 6.3 ± 0.7<sup>***</sup> | 0.14 ± 0.06<sup>***</sup> | 0.69 ± 0.58<sup>***</sup> | 97.6 | 4.76 ± 2.88<sup>***</sup> |

TPA: texture profile analysis, *n* = 9–11 per product; CANN: constitutive artificial neural network, *μ* = 2 ∂*ψ*/∂*I*<sub>1</sub>\|<sub>*I*<sub>1</sub>=3</sub> of the discovered model; we train each model on the mean tension, compression, and shear curves simultaneously and report no dispersion; Sensory: consumer testing, *n* = 100–105 per product; all values are mean ± SD; markers beside TPA values refer to Welch's *t*-tests and markers beside liking to paired Wilcoxon signed-rank tests with Holm correction, each against the dairy version of the same cheese: \*\* *p* < 0.01, \*\*\* *p* < 0.001.

The force–time curves reveal additional differences in compression behavior (Fig. 3). Plant feta and plant mozzarella reach peak force within the first second of compression and then soften before maximum compression, whereas dairy feta maintains a force plateau. This early peak sets the low TPA hardness of plant mozzarella. Dairy mozzarella retains nearly 80% of its initial peak force during the second compression cycle. Resilience ranges from 0.01 to 0.12 across all six products.

### Mechanical response in tension, compression, and shear

The small-strain response separates the three pairs in a different way from TPA (Table 3; Fig. 5). Plant cheddar has the highest initial stiffness of all six products, about twice that of dairy cheddar in tension and more than three times that in compression and shear. Plant and dairy feta have comparable initial stiffness in all three deformation modes. Plant mozzarella is at least five times stiffer than dairy mozzarella in every mode, the reverse of its lower TPA hardness. Relative standard errors range from 1% to 9% across all measurements. The stress–deformation curves reveal additional differences in nonlinear behavior (Fig. 5). In compression, all six cheeses respond approximately linearly, with mild stiffening or softening at most. In tension, all products except plant mozzarella soften. In shear, plant cheddar and plant feta reach peak stresses at strains of approximately 0.04 and 0.05, respectively, followed by a plateau or decline. At a deformation magnitude of 0.08, tensile and compressive stresses differ by less than 20% for five of the six products. Dairy cheddar is the exception, with a tensile stress approximately three times its compressive stress (27.9 *±* 1.1 vs. 9.3 *±* 0.4 kPa).

**Figure 5:**
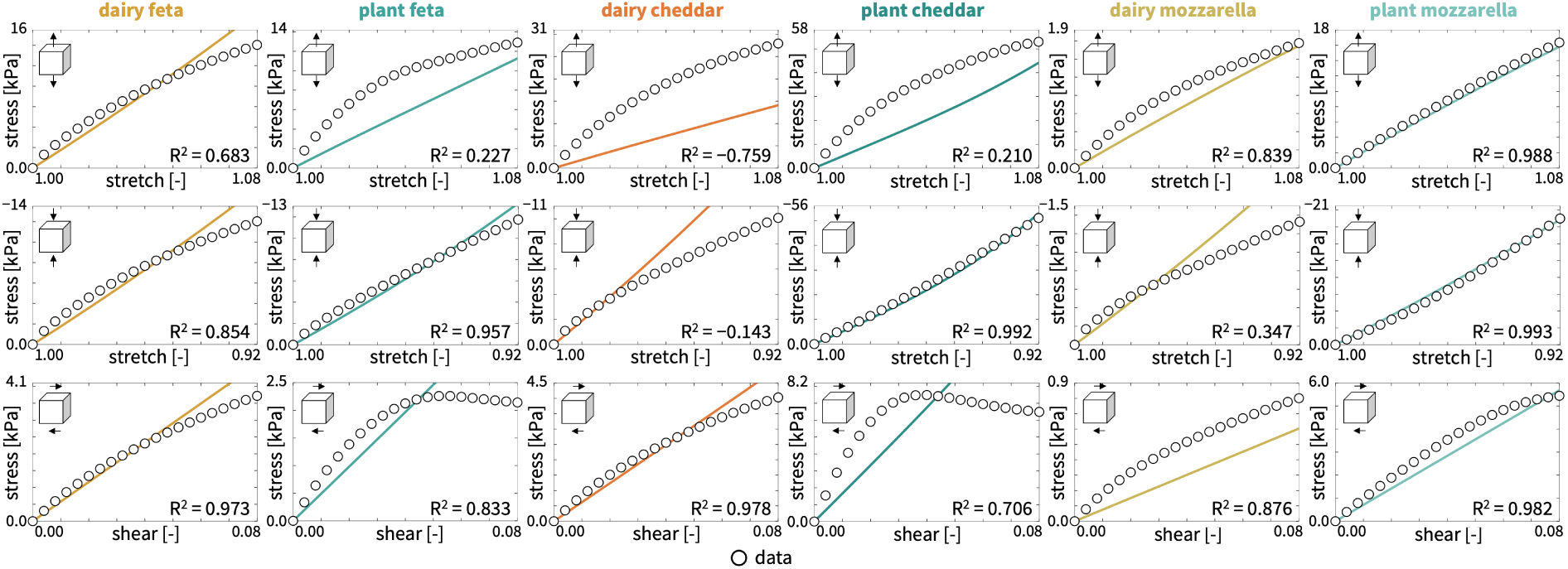
Mechanical response in tension, compression, and shear. Piola stress in tension (top), compression (middle), and shear (bottom) for all six cheese products. Open circles indicate the mean experimental response, and solid lines represent the discovered constitutive models. Annotations report the coefficient of determination *R*^2^ for each deformation mode.

**Table 3:** Initial stiffness in tension, compression, and shear.

| **Cheese** | **Tension**<br>(kPa) | **Compression**<br>(kPa) | **Shear**<br>(kPa) |
|---|---|---|---|
| *Cheddar* |  |  |  |
| Dairy | 566 ± 22 (*n* = 10) | 156 ± 8 (*n* = 14) | 79 ± 6 (*n* = 13) |
| Plant | 1171 ± 24 (*n* = 10) | 556 ± 24 (*n* = 14) | 281 ± 7 (*n* = 11) |
| *Feta* |  |  |  |
| Dairy | 288 ± 22 (*n* = 10) | 223 ± 15 (*n* = 12) | 74 ± 3 (*n* = 13) |
| Plant | 323 ± 11 (*n* = 10) | 187 ± 7 (*n* = 14) | 67 ± 3 (*n* = 14) |
| *Mozzarella* |  |  |  |
| Dairy | 35 ± 3 (*n* = 10) | 25 ± 1 (*n* = 13) | 17 ± 1 (*n* = 14) |
| Plant | 230 ± 6 (*n* = 10) | 193 ± 10 (*n* = 13) | 93 ± 1 (*n* = 14) |

Initial stiffness is the slope of the stress–deformation curve over strain magnitudes from 0.00 to 0.02; values represent mean ± SEM in kPa; parentheses give the number of specimens for each deformation mode.

### Discovered models and fits

The best-in-class one- and two-term models show similar error patterns for all six cheeses (Fig. 6). Models based on the first invariant *I*_1_ consistently outperform models based on the second invariant *I*_2_, and the linear and exponential terms in *I*_1_ give the lowest errors. Dairy and plant-based cheeses therefore share a common family of constitutive models, even though the selected model and its parameters differ within each pair.

**Figure 6:**
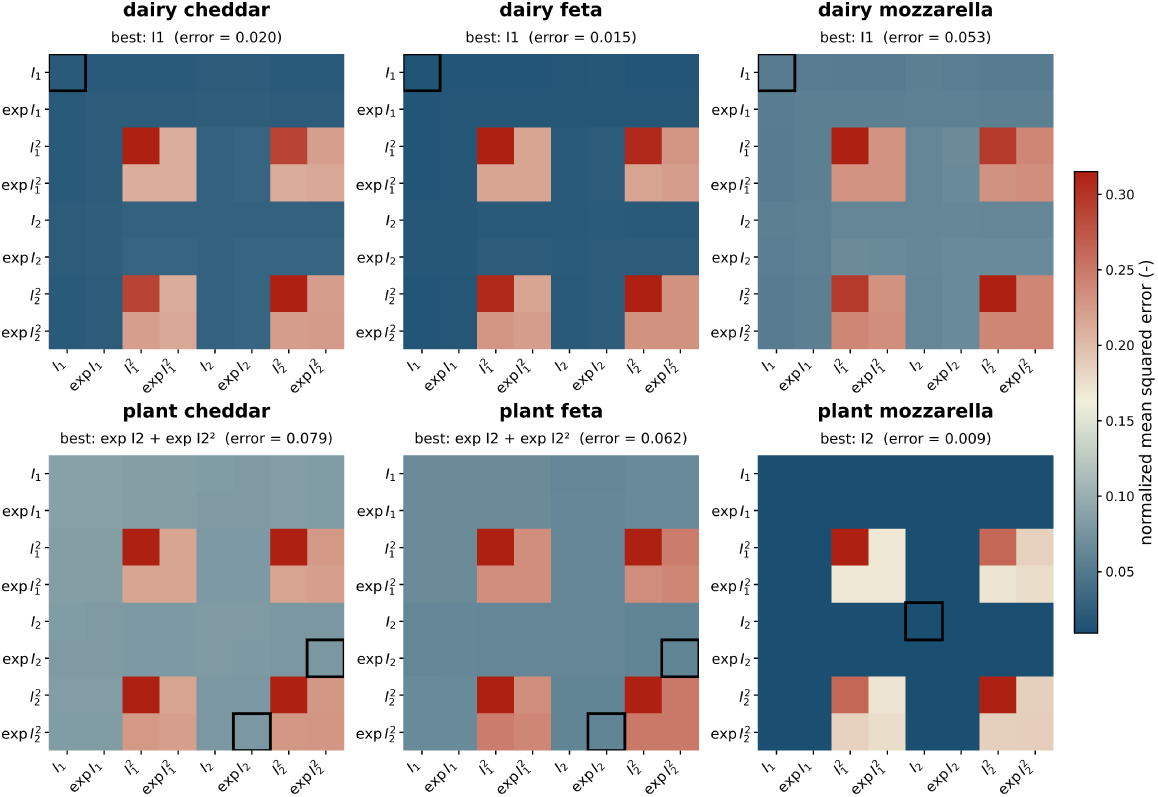
Best-in-class models for dairy and plant-based cheeses. Normalized mean squared error for one-term models (diagonal) and two-term models (off-diagonal) for dairy cheddar, feta, and mozzarella (top) and their plant-based counterparts (bottom). The eight candidate terms are the linear, exponential linear, quadratic, and exponential quadratic terms of the first and second invariants. Colors indicate model error, from dark blue (lowest) to dark red (highest). All six products show the lowest errors for the linear and exponential terms of the first invariant.

Figure 7 shows the discovered models, their fits, and the stress contribution of each term. All discovered models depend on *I*_1_ alone, raised to the first power, with the identity activation, the exponential activation, or both. Model discovery selects the neo-Hookean model for dairy feta and plant mozzarella,

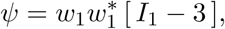

with *w*_1_*w*_1_*^∗^* = 29.11 kPa for dairy feta and *w*_1_*w*_1_*^∗^* = 35.52 kPa for plant mozzarella. For dairy cheddar, dairy mozzarella, and plant feta, it selects the neo-Hookean model plus an exponential term,

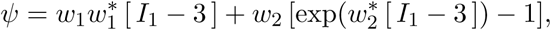

with weights *w*_1_*w*_1_*^∗^* = 18.85 kPa, *w*_2_ = 3.949 kPa, and *w*_2_*^∗^* = 3.069 for dairy cheddar, *w*_1_*w*_2_*^∗^* = 3.027 kPa, *w*_2_ = 1.108 kPa, and *w*_1_*^∗^* = 0.670 for dairy mozzarella, and *w*_1_*w*_2_*^∗^* = 12.19 kPa, *w*_2_ = 3.687 kPa, and *w*_2_*^∗^* = 3.344 for plant feta. For plant cheddar, it selects a single exponential Demiray-type model,

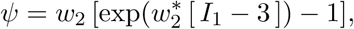

with *w*_2_ = 8.490 kPa and *w*_2_*^∗^* = 9.778. By mean *R*^2^ across the three modes, the discovered model fits plant mozzarella best (0.99), followed by dairy feta (0.83), dairy mozzarella (0.69), plant feta (0.67), plant cheddar (0.64), and dairy cheddar (0.03; tension *R*^2^ = *−*0.76).

**Figure 7:**
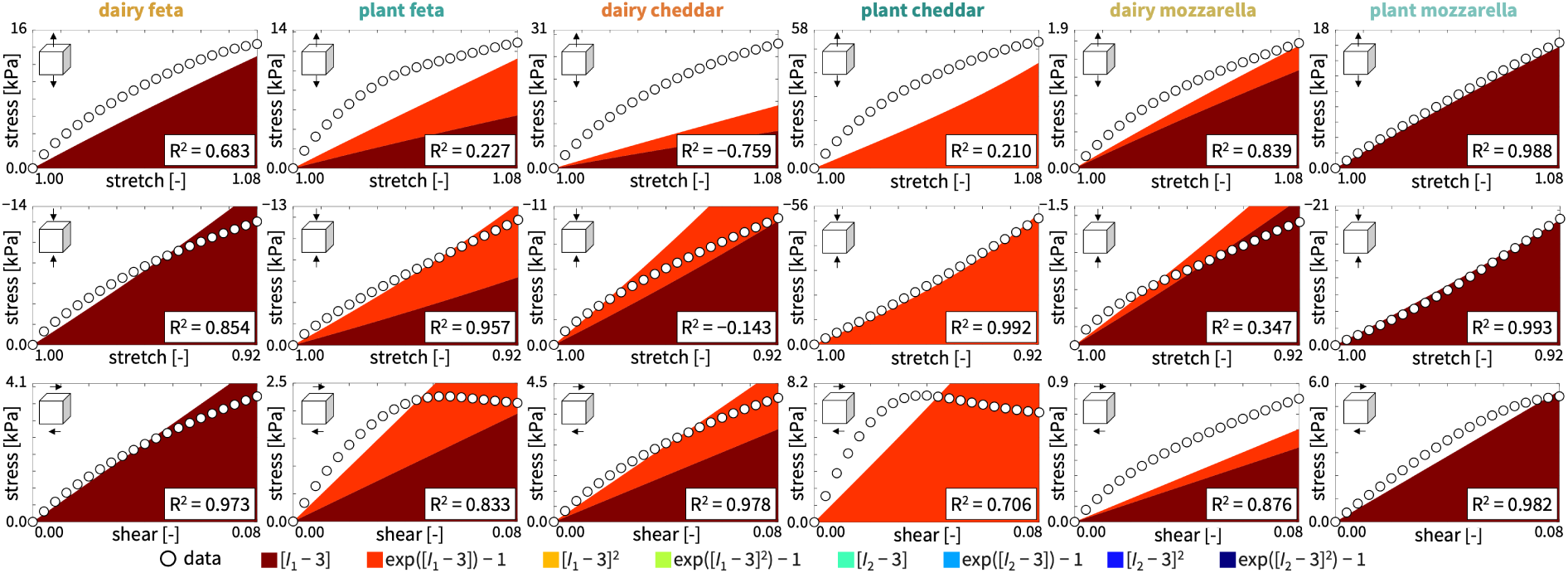
Discovered models and stress contributions. Piola stress in tension (top), compression (middle), and shear (bottom) for all six cheese products. Colored areas indicate the stress contributions of individual model terms, with colors corresponding to Fig. 2. Open circles indicate the mean experimental response. Annotations report the coefficient of determination *R*^2^ for each deformation mode.

### Consumer sensory evaluation

The consumer panel includes 306 participants, and 50.3% of them are between 18 and 30 years old. Participants identify as male (49.3%), female (44.1%), or non-binary or gender non-conforming (6.5%), and predominantly as White (57.8%) or Asian (32.4%). Most participants report omnivorous (65.0%) or flexitarian (20.3%) diets, followed by vegetarian (8.2%) and pescatarian (6.5%) diets. Most eat cheese sometimes (45.5%) or often (30.2%).

#### Overall liking

Consumers consistently prefer the dairy cheeses over their plant-based counterparts (Table 2). Dairy cheeses receive higher overall liking scores for cheddar (7.17 vs. 3.82), feta (6.97 vs. 5.36), and mozzarella (7.61 vs. 4.76; all *p*_adj_ *<* 0.001). Presentation order has no significant effect on the dairy–plant difference in overall liking (*p*_adj_ = 1.00).

#### Sensory attributes

Dairy flavor and tastiness show the largest differences between dairy and plant-based cheeses. On the 0–100 visual analog scale, dairy mozzarella and dairy cheddar receive higher ratings than their plant-based counterparts for dairy flavor (+51.1 and +44.1 points, respectively) and tastiness (+38.7 and +42.9 points, respectively). Plant mozzarella and plant cheddar receive higher graininess ratings (+21.5 and +14.9 points, respectively). All six differences remain significant after Holm correction (*p*_adj_ *<* 0.001).

#### Just-about-right ratings and penalties

Dairy cheeses receive substantially more “just about right” ratings than their plant-based counter-parts (Fig. 8): 78% vs. 29% for cheddar, 74% vs. 38% for feta, and 68% vs. 36% for mozzarella. The largest mean liking penalties occur for excessive sharpness in plant feta (4.33 points), excessive moisture in plant feta (2.84 points), excessive firmness in plant feta (2.81 points), excessive sharpness in plant mozzarella (2.77 points), and insufficient sharpness in plant cheddar (2.70 points).

**Figure 8:**
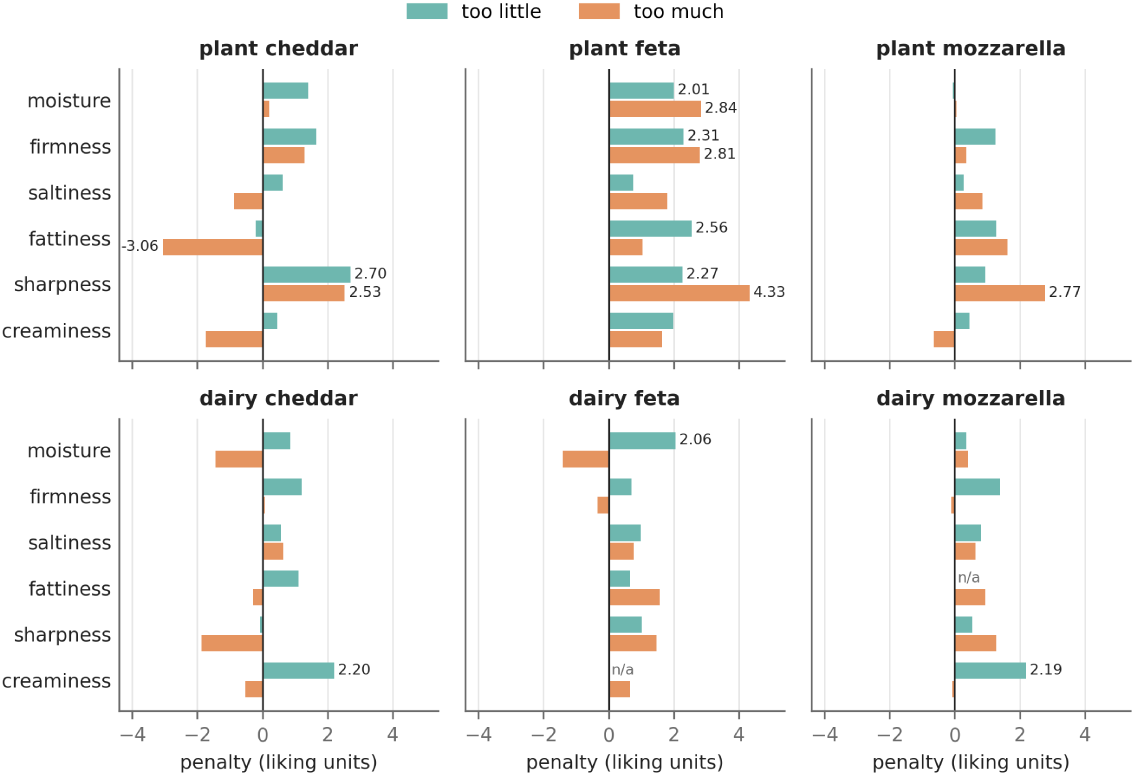
Just-about-right penalty analysis. Mean liking penalties relative to “just about right” ratings for “too little” (teal) and “too much” (orange) ratings of six attributes. Positive values indicate lower mean liking than in the corresponding “just about right” group. Labels identify penalties of 2 points or greater. n/a marks categories with no respondents.

### Instrumental–sensory correlations

Instrumental texture parameters correlate positively with perceived chewiness across the six cheese products (Fig. 9). TPA chewiness (*r* = 0.87, unadjusted *p* = 0.026) and cohesiveness (*r* = 0.94, unadjusted *p* = 0.006) correlate positively with sensory chewiness. TPA hardness shows weaker associations with sensory hardness (*r* = 0.68, unadjusted *p* = 0.14) and softness (*r* = *−*0.69, unadjusted *p* = 0.13). None of these correlations remain statistically significant after Holm correction across all 48 comparisons. Plant mozzarella is the main exception to these trends: consumers perceive it as relatively hard and chewy despite its low TPA hardness and chewiness.

**Figure 9:**
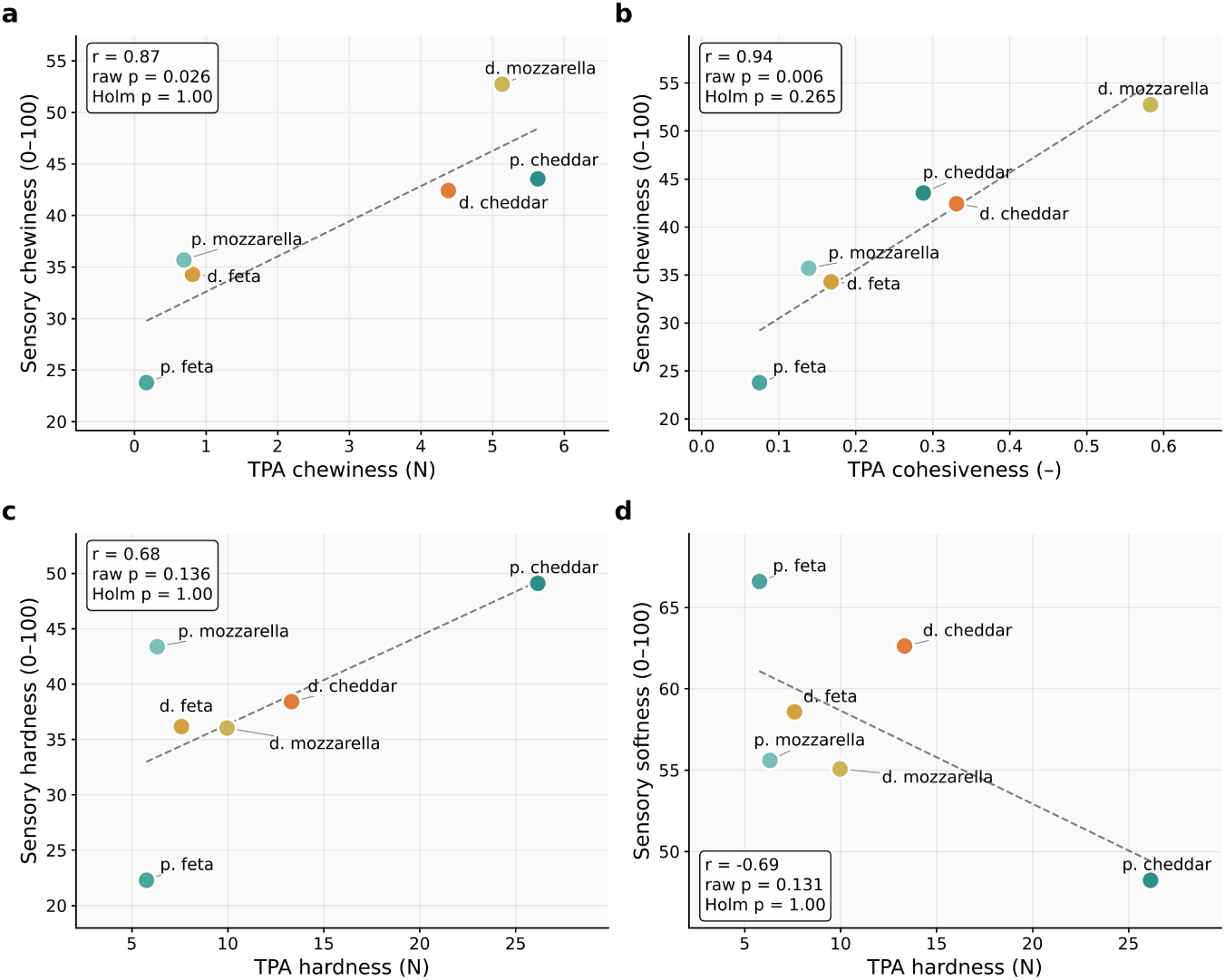
Instrumental–sensory correlations. Pearson correlations between mean texture profile analysis parameters and mean sensory ratings across the six products. Annotations report unadjusted *p*-values. None of the correlations remain statistically significant after Holm correction across 48 comparisons.

## Discussion

Across the six products, dairy cheeses receive higher overall liking scores and more “just about right” ratings than their plant-based counterparts. Automated model discovery selects only linear and exponential terms in the first invariant for all six cheeses, but the selected models differ between the dairy and plant-based product of each style. Instrumental chewiness and cohesiveness correlate with perceived chewiness, although neither association remains significant after correction for multiple comparisons. Deviations in sharpness produce the largest JAR penalties. Because each group contains only one product per cheese style, these findings characterize the tested products rather than dairy and plant-based cheeses as general categories.

### Mechanical signatures differ across cheese products

TPA reveals large, product-specific differences. Plant cheddar has higher hardness than dairy cheddar, whereas plant feta and plant mozzarella have lower hardness and cohesiveness than their dairy counterparts. Plant feta and plant mozzarella have cohesiveness values of 0.08 and 0.14, compared with 0.17–0.58 for the dairy cheeses. Small-strain tests reverse the ranking for mozzarella: plant mozzarella is at least five times stiffer than dairy mozzarella in tension, compression, and shear. TPA hardness reflects the peak force under 50% compression, and plant mozzarella reaches a low peak force early in compression and then softens (Fig. 3). Plant mozzarella therefore resists small deformations more than dairy mozzarella but yields at a lower force under large deformation. Previous work also identifies differences in structural cohesion and failure behavior between dairy and plant-based cheddar and mozzarella, although consumers do not perceive all instrumental differences in cream cheese ^15^.

### No model term is specific to dairy or plant-based cheese

The discovered constitutive models differ between the dairy and plant-based products within each cheese style but show no consistent distinction between the two groups (see the section on discovered models and fits). No model term occurs exclusively in dairy or plant-based cheeses, and the best-in-class analysis places all six cheeses in the same low-error region of linear and exponential *I*_1_ terms (Fig. 6). In meat, all animal products require a quadratic or exponential term, whereas most plant-based products do not^24^. This distinction does not extend to the cheeses in our study. The convex architecture of our network also prevents the discovered models from reproducing the shear peaks of plant cheddar and plant feta. At a deformation magnitude of 0.08, plant feta most closely matches its dairy counterpart across all three deformation modes. Plant cheddar and plant mozzarella sustain substantially higher stresses than their dairy counterparts. Plant mozzarella shows the largest differences, with stresses nearly an order of magnitude higher than those of dairy mozzarella in every mode. Plant cheddar and plant feta show shear peaks that none of the dairy products show. Both products contain starch and oil as principal ingredients (Table 1). Plant mozzarella, the only cashew-based product, responds approximately linearly in all three modes and shares the neo-Hookean model with dairy feta. Our study cannot establish whether the base ingredients cause these differences. Such an analysis requires controlled formulations with systematic variations in starch, fat, and protein content. These results reveal three distinct ways in which plant-based cheeses deviate from their dairy counterparts. Plant mozzarella differs in stiffness, plant feta matches the initial stiffness but differs in its shear response, and plant cheddar differs in both. Formulation targets must therefore account for both initial stiffness and deformation-dependent behavior. Controlled formulation datasets can then translate these mechanical targets into specific recipes.

### Cheese style determines the formulation target

Plant-based cheeses lack a common structural base comparable to milk, and the three plant-based products in our study differ in formulation, mechanics, and consumer response. No single formulation adjustment can therefore close the gap across all three cheese styles. An increase in stiffness may benefit a product that is too soft or weakly cohesive but may worsen an already firm product such as plant cheddar. Equal TPA hardness also does not guarantee equal shear, fracture, melt, or oral behavior. Plant cheddar differs from dairy cheddar in its higher hardness and graininess. Plant feta shows penalties for moisture, firmness, and sharpness in both directions, which suggests that consumers disagree on the direction of these deviations rather than their presence. Plant mozzarella combines high small-strain stiffness with low cohesiveness under large compression and weak dairy flavor. These findings identify formulation targets for the individual products rather than general requirements for each cheese style.

### Instrumental texture captures part of perceived texture

Instrumental measurements and oral perception probe different deformation regimes. The constitutive models characterize mechanical responses up to 8% deformation, TPA compresses specimens to 50%, and mastication fractures the cheese into smaller particles. The discovered models therefore describe the small-deformation response before the first bite, and our study does not establish a direct relationship between constitutive model form and sensory perception. Such a relationship remains a hypothesis that requires mechanical tests at larger deformations, including failure, and consumer evaluations across more products. Across the six products, TPA chewiness and cohesiveness correlate positively with perceived chewiness (*r* = 0.87 and *r* = 0.94), but neither association remains significant after correction for multiple comparisons (*n* = 6). Plant mozzarella deviates most strongly from these trends, with low instrumental hardness and chewiness but relatively high perceived hardness and chewiness. Its high small-strain stiffness may contribute to this mismatch, because the first bite probes the initial resistance of the sample before it fractures. Different mechanical tests capture different aspects of oral texture, and their relevance depends on the cheese style and sensory attribute. Large-amplitude oscillatory shear, for example, characterizes nonlinear energy storage and dissipation beyond the scope of TPA. Previous work links one such parameter to perceived creaminess, fattiness, and moistness in dairy and plant-based cheddar, mozzarella, and cream cheese ^15^.

### Flavor remains a major barrier

Dairy cheeses receive higher overall liking scores in all three matched comparisons (7.17 vs. 3.82 for cheddar, 6.97 vs. 5.36 for feta, and 7.61 vs. 4.76 for mozzarella) and substantially more “just about right” ratings (68–78% vs. 29–38%). These findings align with the low retail adoption of plant-based cheese ^4;8^. Excessive sharpness produces the largest JAR penalties in plant feta and plant mozzarella. These penalties suggest that flavor contributes to lower acceptance, but they do not provide chemical evidence of specific off-flavors. Participants may interpret sharpness as acidity, pungency, an undesirable plant-derived note, or a combination of these sensations. Previous studies also identify artificial flavors, off-flavors, and gritty or rubbery textures as barriers to consumer acceptance of plant-based cheddar and mozzarella^8;36^. Flavor and texture therefore both limit acceptance. Plant-based cheese does not always receive lower consumer acceptance than dairy cheese. In a study of cream cheese, the largest consumer segment rated two plant-based and two dairy products equally, with creamy and smooth mouthfeel as the main drivers of acceptance ^37^. Plant-based barista milk and creamer have also reached substantial market shares^8^. Cheese, however, must satisfy a wider range of culinary uses than these products.

### Limitations and future directions

Our study has several limitations. First, each cheese style includes only one dairy and one plant-based product, which confounds brand and formulation with the dairy–plant contrast. Commercial formulations may also change over time. Broader conclusions require more products or controlled formulations per style. Second, each participant evaluates only one cheese style, so cross-style comparisons involve different panels, whereas dairy–plant comparisons remain paired. Visible differences may influence ratings despite three-digit codes (Table 1). All tests use unheated samples at room temperature, although consumers often heat mozzarella and cheddar before they eat them. Future studies should examine temperature-dependent rheology, melt, oil and water release, and stretch. Third, the constitutive models cover deformations of at most 8%, far below those of mastication (see the section on instrumental and perceived texture). Plant cheddar and plant feta show shear peaks near strains of 0.04–0.05, unlike the other products under the same protocol, which makes a systematic fixture artifact less likely. The convex network cannot capture this softening, which contributes to poor fits. Models with damage or viscoelasticity may capture this response. Fourth, the models assume isotropic incompressibility, which requires the initial uniaxial stiffness to equal three times the initial shear stiffness. Experimental ratios range from 1.5 to 3.0 in compression and from 2.1 to 7.2 in tension (Table 3). These deviations may reflect material behavior, experimental differences, or both. Finally, tensile tests use different instruments, specimen geometries, and grips than compression and shear tests. At a deformation of 0.08, five cheeses show similar tensile and compressive stresses, whereas dairy cheddar sustains three times higher stress in tension. This difference may reflect material behavior or experimental conditions. A single isotropic model cannot reproduce this tension–compression asymmetry, which accounts for the poor fit for dairy cheddar (mean *R*^2^ = 0.03; tension *R*^2^ = *−*0.76).

In summary, the plant-based cheeses in this study differ from their dairy counterparts in product-specific ways. Consumers rate the plant-based products 3.8–5.4 on overall liking, compared with 7.0–7.6 for the dairy products. Plant mozzarella differs in stiffness, plant feta in its nonlinear shear response, and plant cheddar in both, and no model term separates dairy from plant-based cheese. Excessive sharpness produces the largest sensory penalties, so formulation must address flavor alongside texture. Studies with more products per style can test whether these findings hold beyond the six products tested here.

## Methods

### Cheese products

We test one plant-based and one dairy product for each of three cheese styles, cheddar, feta, and mozzarella (Table 1). We purchase all products from local retailers in Stanford, California, store them at 4 *^◦^*C, and test them at room temperature within their labeled shelf-life dates.

### Texture profile analysis

We perform TPA on a TA Instruments Discovery HR-2 rheometer in axial compression with a flat stainless steel probe of 8 mm diameter^18^. We cut cylindrical specimens from the cheese blocks with an 8 mm stainless steel punch and equilibrate them at a room temperature of 22 *±* 1 *^◦^*C for 30 min before each test. We test *n* = 10–12 specimens per product. We define the initial specimen height as the probe gap at the start of the test. Heights differ among products: 8.2 *±* 0.1 mm and 7.3 *±* 0.6 mm for plant and dairy cheddar, 9.8 *±* 0.1 mm and 12.0 *±* 0.1 mm for plant and dairy feta, and 10.4 *±* 0.1 mm and 10.4 *±* 0.1 mm for plant and dairy mozzarella.

We establish initial probe contact at forces of 0.0–2.2 N and compress each specimen twice to 50% of its initial height. Each compression reaches the maximum displacement in 2.3 s, which corresponds to a nominal strain rate of approximately 0.22/s and a probe speed of 1.7–2.7 mm/s, depending on specimen height. Between cycles, the probe returns to its initial position and immediately begins the second compression, with no dwell time^19^. We record force and gap at 244 Hz and smooth the force signal with a Gaussian filter with *σ* = 2 data points.

All specimens reach peak force before maximum compression. We therefore separate the loading and unloading portions of each cycle at the minimum gap rather than at peak force. We calculate the magnitudes of loading and unloading work from the force–displacement curves and exclude negative forces from adhesion. We denote the first-cycle loading and unloading work by *A*_1_ and *A*_2_, and the second-cycle values by *A*_3_ and *A*_4_.

We extract six parameters from each test^18;19^. The stiffness index is the peak nominal stress during the first compression divided by the prescribed maximum strain, *E*_index_ = (*F*_1_*/A*)*/*0.5, where *A* = 50.27 mm^2^ is the initial specimen cross-sectional area. Hardness is the peak force during the first compression cycle, *F*_1_. Cohesiveness is the ratio of the total positive-force work magnitudes in the second and first cycles, (*A*_3_ +*A*_4_)*/*(*A*_1_ +*A*_2_). Springiness is the ratio of the second- to first-cycle probe travel from contact to maximum compression, *d*_2_*/d*_1_. Resilience is the ratio of unloading to loading work in the first cycle, *A*_2_*/A*_1_. Chewiness is the product of hardness, cohesiveness, and springiness, *F*_1_[(*A*_3_ + *A*_4_)*/*(*A*_1_ + *A*_2_)](*d*_2_*/d*_1_). The stiffness index differs from the classical TPA stiffness because peak force precedes maximum compression in our tests.

### Compression and simple shear testing

We measure compression and simple shear on the same rheometer with an 8 mm parallel plate geometry fitted with 3D-printed sandpaper surfaces to prevent slip (Fig. 1a). We cut cylindrical specimens of 8 mm diameter and 2 mm height from separate cheese blocks and allow them to equilibrate for 5 min after placement between the plates. We apply both deformation modes monotonically at a constant rate of 0.002/s and record nominal stress against applied strain. Across all six products, we test *n* = 80 specimens in compression and *n* = 79 in shear. We report compression stress as a positive magnitude and convert it to signed nominal stress for the constitutive equations.

### Tensile testing

We adapt a uniaxial tensile protocol^24^. We cut rectangular specimens of 10 mm width, 10 mm thickness, and 40 mm length from each cheese block and secure them in Instron 2710-103 screw side-action grips with 1 kN capacity, fitted with 3D-printed serrated faces (Fig. 1b). We mount the grips on an Instron 5848 frame (Instron, Canton, MA) with a 100 N load cell and set the gauge length to approximately 20 mm. We apply a preload of approximately 0.5 N to remove slack before we record the initial gauge length. We stretch the specimens quasi-statically at a stretch rate of 0.002/s until failure, which we identify as a marked drop in stress. We test *n* = 60 specimens intension, 10 per product, at a room temperature of 22 *±* 1 *^◦^*C. We convert force and displacement to nominal stress and stretch, smooth the resulting curves, and resample each curve to 500 points.

### Data processing

We restrict all three deformation modes to a common deformation magnitude of 0.08: tensile stretch from 1.00 to 1.08, compressive stretch from 0.92 to 1.00, and shear strain from 0.00 to 0.08. We interpolate each curve onto a common 81-point grid and average the curves for each product and mode. We report the standard error of the mean (SEM) across replicates. Initial stiffness is the slope of a linear fit to each curve over deformation magnitudes from 0.00 to 0.02, which we summarize as the mean *±* SEM.

### Continuum mechanics

The motion ***x*** = ***φ***(***X***) maps the undeformed configuration, ***X***, to the deformed configuration, ***x***. The deformation gradient is ***F*** = *∂****x****/∂****X***, and the right Cauchy–Green tensor is ***C*** = ***F*** ^t^ *·* ***F*** . We model each cheese as an isotropic, incompressible, hyperelastic solid with *I*_3_ = det(***C***) = *J* ^2^ = 1, where *J* = det(***F*** ). The strain energy function *ψ* therefore depends only on the first and second invariants of the right Cauchy–Green tensor, *I*_1_ = tr (***C***) and *I*_2_ = <u>^1^</u> [(tr ***C***)^2^ *−* tr(***C***^2^)] ^20^. The first Piola–Kirchhoff stress of an incompressible hyperelastic solid, ***P*** = *∂ψ/∂****F*** *− p* ***F*** *^−^*^t^, relates the force in the current configuration to the area in the undeformed configuration through the strain energy derivative, *∂ψ/∂****F*** , and the pressure constraint, *−p* ***F*** *^−^*^t^, where the Lagrange multiplier *p* enforces incompressibility. For uniaxial tension or compression with stretch *λ*, the invariants reduce to *I*_1_ = *λ*^2^ + 2*/λ* and *I*_2_ = 2*λ* + 1*/λ*^2^, and the nominal axial stress is *P*_11_ = 2 (*λ −* 1*/λ*^2^) [*∂ψ/∂I*_1_ + (1*/λ*)*∂ψ/∂I*_2_]. For simple shear with shear strain *γ*, the invariants are *I*_1_ = *I*_2_ = 3 + *γ*^2^, and the nominal shear stress is *P*_12_ = 2*γ* [*∂ψ/∂I*_1_ + *∂ψ/∂I*_2_].

### Constitutive artificial neural network

Constitutive artificial neural networks automate model discovery and satisfy thermodynamic consistency, material objectivity, material symmetry, and polyconvexity by design^20^. We adopt the eight-term architecture from previous studies of meat ^23;24^. The network takes the invariants *I*_1_ and *I*_2_ as input. Its first hidden layer raises [*I*_1_ *−* 3] and [*I*_2_ *−* 3] to the first and second powers, and its second hidden layer applies the identity (*◦*) and the exponential exp(*◦*) *−* 1. Each of the *n* = 8 terms has an internal weight *w_i_^∗^* and an external weight *w_i_*, for a total of 16 weights (Fig. 2). The strain energy function is

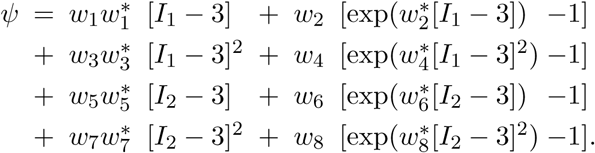

We constrain all weights to non-negative values, so each term is convex and monotonically increasing over the admissible invariant range. These constraints ensure polyconvexity of the strain energy function. The first Piola–Kirchhoff stress follows as 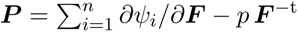, where *ψ_i_* denotes the contribution of term *i* to the strain energy and *p* enforces incompressibility.

### Network training and model selection

We train the network simultaneously on the mean tension, compression, and shear curves of each cheese, with 41 points per curve. We define the loss,

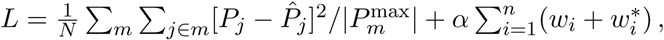

as the mean squared error normalized by the absolute peak stress of each mode, plus an *L*_1_ regularization that controls the number of active terms. Here, *m* denotes the deformation mode, *P_j_* the predicted stress, *P*^^^*j* the measured mean stress, *P_m_*^max^ the peak stress of mode *m*, *N* the number of grid points per output, *n* = 8 the number of terms, and *α* the regularization parameter. We minimize the loss with the Adam optimizer at a learning rate of 0.001 and a batch size of 8 for at most 20,000 epochs. We stop training early if the loss shows no improvement for 300 consecutive epochs.

For model discovery, we sweep *α* over 0, 10*^−^*^4^, 10*^−^*^3^, 10*^−^*^2^, and 10*^−^*^1^. We train eight random initializations at *α* = 0 and four at each other value, and we retain the initialization with the lowest training loss at each *α*. The fit at *α* = 0 serves as the reference. For each cheese, we select the largest *α* for which the *R*^2^ of every deformation mode stays within 0.05 of the reference. This criterion gives *α* = 10*^−^*^3^ for dairy cheddar and dairy mozzarella, *α* = 10*^−^*^2^ for dairy feta and plant feta, and *α* = 10*^−^*^1^ for plant cheddar and plant mozzarella. We report model quality as the coefficient of determination *R*^2^ between the mean measured curve and the model.

To map model error across candidate models, we also train all 8 one-term and all 28 two-term models of the eight candidate terms for each cheese without regularization, and we record the normalized mean squared error of each model^21^.

### Consumer panel

Between May and June 2026, we recruit *n* = 306 participants from the Stanford University community through email lists and posted flyers. Eligible participants are at least 18 years old, can read English, and are willing to eat both dairy and plant-based cheese. We exclude individuals who report allergies to milk, soy, pea protein, coconut, nuts, wheat, or legumes, or disorders that affect taste or smell. Participants provide electronic informed consent through Qualtrics and receive a $5 gift card. The Stanford University IRB approves the study under eProtocol #85843.

### Sensory procedure

We conduct sensory evaluations at individual table stations in a university dining hall. Each participant evaluates the plant-based and dairy versions of one cheese style: *n* = 105 evaluate cheddar, *n* = 101 evaluate feta, and *n* = 100 evaluate mozzarella. We cut samples into 1 cm^3^ cubes and serve them at a room temperature of 22 *^◦^*C within 2 h of preparation. We label samples with random three-digit codes and randomize presentation order across participants: 133 participants receive the plant-based sample first, 172 receive the dairy sample first, and one participant lacks presentation-order data. We provide filtered water and unsalted crackers as palate cleansers.

### Sensory attribute ratings

Participants rate overall liking on a nine-point scale from 1 (dislike very much) to 9 (like very much), with 5 as neutral. They also rate eight texture and flavor attributes on 0–100 visual analog scales: chewiness, hardness, moisture, graininess, dairy flavor, fattiness, tastiness, and softness. The chewiness scale ranges from 0 (breaks down instantly) to 100 (requires repeated chews), and the hardness scale ranges from 0 (extremely soft) to 100 (very hard).

### Just-about-right scales

Participants rate six attributes, moisture, firmness, saltiness, fattiness, sharpness, and creaminess, on a five-point just-about-right (JAR) scale, where 1 denotes much too little, 3 just about right, and 5 much too much ^38^.

### Statistical analysis

We perform all analyses in Python 3.11 with scipy (v1.11) and statsmodels (v0.14). For TPA, we exclude a specimen if any parameter falls outside the middle 50% of values for that product by more than 1.5 times the span of that range, or differs from the product mean by more than 2.5 standard deviations. This rule leaves *n* = 9–11 specimens per product. We use Welch’s *t*-tests to compare the plant-based and dairy versions of each cheese and Tukey’s HSD to compare all six cheeses. For sensory data, we use paired Wilcoxon signed-rank tests to compare plant-based and dairy ratings within each cheese style. We apply the Holm correction to control the family-wise error rate across the 27 liking and visual analog scale comparisons. We test presentation-order effects on each participant’s dairy-minus-plant contrast with 10,000-permutation tests that stratify permutations by cheese style, and we apply the Holm correction across nine outcomes. For JAR penalty analysis, we subtract the mean liking score in each deviation category (too little or too much) from the mean liking score in the just-about-right category; a positive penalty indicates lower liking in the deviation category38. We calculate Pearson correlations between mean TPA parameters and mean sensory ratings across the six cheeses (*n* = 6) and apply the Holm correction across the 48 correlations. We treat these aggregate correlations as exploratory. We set the significance threshold at *α* = 0.05. For sensory comparisons, presentation-order tests, and TPA–sensory correlations, significance markers refer to Holm-adjusted *p*-values: *^∗∗∗^* for *p*_adj_ *<* 0.001, *^∗∗^* for *p*_adj_ *<* 0.01, *^∗^* for *p*_adj_ *<* 0.05, and ns for not significant.

### Use of generative artificial intelligence

We used Claude (Anthropic) to help write analysis code and manuscript text. All authors reviewed and edited the code and text, and they take full responsibility for the content of this article.

## Data availability

Tension, compression, and shear data and de-identified sensory data will be available at https://github.com/LivingMatterLab/AI4Food.

## Code availability

Texture profile analysis processing scripts and constitutive neural network code will be available at https://github.com/LivingMatterLab/AI4Food. The constitutive neural network implementation follows https://github.com/LivingMatterLab/CANN.

## Author contributions

A.O.K.: Conceptualization, Software, Formal analysis, Visualization, Writing – original draft, Funding acquisition. E.C.D.: Investigation, Software, Formal analysis, Visualization. V.E.: Investigation, Software, Formal analysis, Visualization. E.K.: Software, Formal analysis, Visualization, Supervision, Project administration, Funding acquisition. All authors reviewed and approved the final manuscript.

